# Invasion pathway predicts the axis of ecological niche reorganisation in freshwater crayfish

**DOI:** 10.64898/2026.04.05.716527

**Authors:** Kristian miok, Olga n. Petko, Marko robnik-šikonja, Lucian pârvulescu

## Abstract

**Aim:** Understanding whether invasive species retain or shift their ecological niches has traditionally relied on scalar overlap metrics that quantify the magnitude of niche change, but not its structure. Here, we test whether biological invasions involve a reorganisation of the environmental axes along which native and invasive ranges are differentiated, and whether the dominant axes of this reorganisation are consistently associated with invasion pathway type (intercontinental vs. within-continent).

**Location:** Global (North America, Europe, Africa, Asia, Australasia).

**Time period:** Contemporary (environmental variables representing long-term averages, 1980–2021).

**Major taxa studied:** Freshwater crayfish (Decapoda: Astacidea): *Procambarus clarkii*, *Faxonius limosus*, *Pacifastacus leniusculus*, *Faxonius virilis*, *Faxonius rusticus*.

**Methods:** We analysed native and invasive occurrences for five globally important crayfish invaders using ∼400 hydrologically resolved environmental variables from the Global Crayfish Database of Geospatial Traits. Classification models were used to quantify environmental differentiation between native and invasive ranges, and feature contributions were aggregated by environmental domain (climate, topography, soil, land cover). Patterns were evaluated across intercontinental and within-continent invasion pathways and assessed for robustness using cross-validation, permutation tests, sample-size sensitivity, and comparisons with classical niche overlap metrics.

**Results:** Native and invasive occurrences were consistently distinguishable across all species (accuracy 96.5–99.9%). A pathway-dependent pattern emerged: intercontinental invaders were primarily differentiated along climatic dimensions (58–76% of model importance), whereas within-continent invaders showed a more balanced contribution of climatic and topographic variables (∼42% each), including strong signals from river network position. This contrast was stable across cross-validation folds (SD < 1.6%), and supported by permutation tests (P = 0.001). Classical niche overlap metrics (Schoener’s D = 0.30–0.62) did not capture this qualitative distinction.

**Main conclusions:** Biological invasions involve not only changes in niche position but a reorganisation of the environmental axes that distinguish species’ distributions. Our results suggest that the dominant axes of this reorganisation differ systematically with invasion pathway, reflecting whether species encounter novel climatic regimes or primarily shift within existing climatic space along topographic and network-position gradients. By resolving which environmental dimensions underpin native–invasive differentiation, this approach provides a complementary perspective to scalar overlap metrics and a basis for more mechanistic interpretations of invasion processes.

## 1 Introduction

Freshwater ecosystems harbour a disproportionate share of global biodiversity relative to their spatial extent, yet they are among the most imperilled systems on Earth, with freshwater species populations declining at roughly twice the rate observed in terrestrial or marine realms (Dudgeon et al., 2006; Reid et al., 2019; WWF, 2022). Biological invasions rank among the principal drivers of this decline, restructuring recipient communities through competitive displacement, pathogen transmission, and trophic cascading (Strayer & Dudgeon, 2010). Understanding the environmental conditions that enable invasive species to establish and spread is therefore a central challenge for freshwater conservation.

A fundamental question in invasion ecology is whether invasive species occupy the same ecological niche in their invaded range as in their native range, a pattern termed niche conservatism, or whether they undergo niche shifts upon establishment in novel environments (Broennimann et al., 2007; Guisan et al., 2014). Most studies have relied on niche overlap metrics such as Schoener’s D and Warren’s I, which summarise niche similarity as scalar quantities (Petitpierre et al., 2012). While informative, these metrics tell us *how much* niches differ but not *which* environmental axes are reorganised or *why* the reorganisation takes the form it does. A species whose niche shifts along climatic axes because it has crossed an ocean is ecologically distinct from one whose niche shifts along topographic axes because it has colonised a different position within the same river network, yet both may receive similar overlap scores.

Freshwater crayfish (Decapoda: Astacidea) provide an ideal system to investigate this question. They function as keystone consumers and ecosystem engineers, structuring benthic communities, modulating nutrient cycling, and creating habitat heterogeneity through burrowing (Reynolds et al., 2013). Several species rank among the most damaging freshwater invaders worldwide (Oficialdegui et al., 2020; O’Hea Miller et al., 2024), and they include both intercontinental and within-continent invaders, providing a natural comparative framework.

A critical limitation of previous analyses has been their reliance on terrestrial climate grids that ignore the dendritic structure of river networks (Domisch et al., 2015; Cid et al., 2022). The Global Crayfish Database of Geospatial Traits (GeoTraits; Petko et al., in press; data freely accessible at https://world.crayfish.ro/) addresses this by integrating occurrence records from the World of Crayfish® platform (Ion et al., 2024) with ∼400 environmental variables from GeoFRESH (Domisch et al., 2024) at 90 m resolution from Hydrography90m (Amatulli et al., 2022), providing network-aware climatic, topographic, soil, and land-cover descriptors at both local and upstream scales.

Machine learning approaches, particularly decision trees and random forests, identify specific environmental variables and thresholds that distinguish classes, producing interpretable ecological rules (Elith & Leathwick, 2009; Christin et al., 2019). When applied to the native–invasive comparison, they reveal not only whether niches differ but precisely which variables distinguish ranges and at what threshold values.

Here we address three questions: (1) Can native and invasive ranges of globally significant invasive crayfish be reliably distinguished using network-aware environmental variables? (2) Which variable types are most important, and does this differ between intercontinental and within-continent invaders? (3) Do classical overlap metrics capture the same signal as machine learning? We study five species: *Procambarus clarkii*, *Faxonius limosus*, *Pacifastacus leniusculus* (intercontinental) and *Faxonius virilis*, *Faxonius rusticus* (within-continent). Throughout this paper, we use the term ‘niche reorganisation’ to denote changes in the relative importance of environmental axes defining species’ distributions, as distinct from ‘niche shift’, which refers to changes in position within environmental space. We define the ‘axis of niche reorganisation’ operationally as the environmental domain (climate, topography, soil, or land cover) accounting for the largest share of aggregated feature importance in models discriminating native from invasive occurrences.

## 2 Methods

### 2.1 Data source and filtering

Occurrence records and environmental data were obtained from the Global Crayfish Database of Geospatial Traits (GeoTraits; Petko et al., in press), which links 115,191 georeferenced crayfish records from the World of Crayfish® platform (WoC®; Ion et al., 2024) to ∼400 environmental variables extracted through the GeoFRESH framework (Domisch et al., 2024). Environmental variables span four thematic domains: Climate (76 local and 19 upstream variables derived from CHELSA v2.1 BIOCLIM; Karger et al., 2017), Topography (144 local and 40 upstream variables from Hydrography90m; Amatulli et al., 2022), Soil (60 local and 15 upstream variables from SoilGrids250m; Hengl et al., 2017), and Land Cover (22 local and 22 upstream variables from ESA CCI LC). Local variables (prefixed l_) describe conditions at the 90 m stream segment scale; upstream variables (prefixed u_) represent catchment-aggregated means weighted by contributing area.

Records were filtered through a sequential quality pipeline: (1) retention of only ‘Native’ and ‘Alien’ status records, excluding ‘Introduced’ and ‘Type locality’ designations to ensure clear native–invasive contrasts; (2) retention of High spatial accuracy records only, as assigned by data contributors on the basis of georeferencing source precision; (3) exclusion of records with snapping distances >1 km from the nearest Hydrography90m stream segment, as validated in the GeoTraits quality assessment framework; and (4) deduplication to one record per species per status per stream segment (subc_id). This pipeline reduced the dataset from 115,191 to 64,737 records. Species were selected for analysis based on the availability of sufficient records in both native and invasive ranges, with a minimum threshold of 40 records per range context after filtering. Five species met this criterion (Table 1): *Procambarus clarkii* (red swamp crayfish; native to the south-central United States, invasive across Europe, Africa, and Asia), *Faxonius limosus* (spiny-cheek crayfish; native to eastern North America, invasive in Europe), *Pacifastacus leniusculus* (signal crayfish; native to the Pacific Northwest of North America, invasive in Europe), *Faxonius virilis* (virile crayfish; native to central-eastern North America, invasive in western North America and Europe), and *Faxonius rusticus* (rusty crayfish; native to the Ohio River basin, invasive in the Great Lakes region and northeastern United States). These species encompass both intercontinental invasion pathways (*P. clarkii*, *F. limosus*, *P. leniusculus*) and within-continent range expansions (*F. virilis*, *F. rusticus*), providing a natural comparative framework for testing whether the nature of niche reorganisation differs between pathway types.

**Table 1.** Summary of study species and record counts after quality filtering.

| Species | Native (n) | Invasive (n) | Total | Variables | Invasion type |
| --- | --- | --- | --- | --- | --- |
| *P. clarkii* | 2,208 | 8,527 | 10,735 | 195 | Intercontinental |
| *F. limosus* | 382 | 4,089 | 4,471 | 270 | Intercontinental |
| *P. leniusculus* | 117 | 4,459 | 4,576 | 292 | Intercontinental |
| *F. virilis* | 1,856 | 500 | 2,356 | 277 | Within-continent |
| *F. rusticus* | 1,223 | 670 | 1,893 | 271 | Within-continent |

### 2.2 Environmental variable preparation and modelling framework

For each species, the full set of 398 environmental variables was subjected to three species-specific cleaning steps. First, variables with more than 30% missing values across the species’ records were removed. Second, records with more than 50% missing values across retained variables were excluded, and remaining gaps were filled using median imputation. Third, zero-variance variables and one of each pair of variables with Pearson correlation |r| > 0.98 were removed, retaining the variable with lower mean absolute correlation with all others. This species-specific cleaning ensured that models were trained on informative, non-redundant predictor sets, yielding 195–292 variables per species (Table 1).

Two complementary classification approaches were applied to the cleaned datasets. First, for each species, a decision tree classifier was trained to distinguish native-range from invasive-range occurrences using scikit-learn (Pedregosa et al., 2011) with the CART algorithm, class_weight=‘balanced’ to account for unequal class sizes, and a maximum depth of 5. Model performance was evaluated using 5-fold stratified cross-validation, reporting accuracy, F1 score, and area under the receiver operating characteristic curve (ROC AUC).

Depth sensitivity was assessed by training trees at depths 3, 4, 5, and 8, comparing performance and complexity (decision tree visualisations for all species at depth 5 are provided in Figure S9). From each tree, we extracted feature importances (Gini importance), decision rules (root-to-leaf paths), and threshold values at each split node. Feature importances were aggregated by environmental variable type (Climate, Topography, Soil, Land Cover) and spatial scale (Local, Upstream) to characterise the dominant environmental dimensions distinguishing native from invasive ranges.

Second, random forest models were fitted to the same native-versus-invasive datasets to assess whether patterns identified by single decision trees were robust to ensemble classification (Breiman, 2001). For each species, a random forest with 500 trees and balanced class weights was trained using 5-fold stratified cross-validation. Feature importance was quantified in two complementary ways: standard Gini importance extracted from the fitted model, and SHAP values computed using TreeExplainer (Lundberg & Lee, 2017), summarised as mean absolute SHAP value per feature. Theme-level importance was calculated as the sum of feature importances within each environmental domain, enabling direct comparison with decision tree results. At the individual-variable level, the top-ranked features under both Gini and SHAP importance were compared with those identified by the decision trees to assess ecological consistency across methods (Figure S1; species-level SHAP summary plots in Figure S11; complete feature importance rankings in Tables S7–S9).

### 2.3 Classical niche overlap and niche reorganisation analysis

To benchmark the machine learning approach against classical niche overlap summaries, we quantified the overlap between native and invasive environmental space for each species using Schoener’s D and Warren’s I (Warren et al., 2008). For each species, native and invasive records were pooled, and the cleaned environmental variables were standardised. Principal component analysis was then applied to the standardised matrix, and the first two principal components were retained as a reduced representation of environmental space, jointly capturing 20.8–25.0% of total variance across species (Figure S4; Table S5). Although this proportion is modest in absolute terms, it is expected given the high dimensionality of the predictor space (195–292 variables) and is consistent with standard practice in multivariate niche overlap analyses (Broennimann et al., 2007). Smoothed occupancy densities were estimated for native and invasive records separately using binned counts followed by Gaussian smoothing on a regular two-dimensional grid. Each density surface was normalised to sum to one, producing comparable probability distributions. Schoener’s D was calculated as 1 − 0.5∑|p_i_ − q_i_|, and Warren’s I as ∑√(p_i_q_i_), where p_i_ and q_i_ are normalised occupancy probabilities in each grid cell.

To compare the ecological rules governing species occurrence independently in each range, we trained separate presence–background decision trees for the native range and the invasive range. For each range, the focal species’ occurrence records served as presences (target = 1), and an equal number of records from all other species with the same status (Native or Alien) in the filtered dataset served as pseudo-absences (target = 0), providing a realistic environmental background for the corresponding geographic region. Trees were trained with maximum depth 4 and class_weight=‘balanced’, with 5-fold stratified cross-validation. We then compared the top splitting variables and their threshold values between native and invasive niche models. For variables appearing as top splits in both models, we computed the threshold shift (invasive threshold minus native threshold) and its relative magnitude. Variables appearing in only one model were classified as range-specific niche determinants.

### 2.4 Statistical validation and sensitivity analyses

Three additional analyses were conducted to assess the robustness of the main findings. First, a species-level permutation test assessed whether the observed contrast between intercontinental and within-continent invaders in the relative importance of climate versus topography could arise by chance from the assignment of invasion pathway categories. The intercontinental group comprised *P. clarkii*, *F. limosus*, and *P. leniusculus*; the within-continent group comprised *F. virilis* and *F. rusticus*. For each species, the difference between climate and topography importance was calculated from the random forest analysis. The observed dichotomy statistic was defined as the mean species-level value in the intercontinental group minus the mean value in the within-continent group. Group labels were then randomly permuted 999 times while preserving original group sizes (3 intercontinental, 2 within-continent), and the same contrast was recalculated for each permutation. Empirical one-sided and two-sided P-values were computed with a +1 correction. The test was run separately for Gini- and SHAP-based importance, and for both the climate-minus-topography and climate/topography ratio statistics (Table S1; Figure S6).

Variable-importance stability was quantified across cross-validation folds. For each species, a random forest was fitted within each of the five folds, and theme-level importance (climate, topography, soil, land cover) was calculated per fold. Mean and standard deviation across folds were reported for both theme-level and individual-variable importance, allowing us to assess whether the broad dichotomy and specific driver identities were stable across data partitions (Table S2; Figures S2–S3).

Sample-size sensitivity analysis tested whether the smaller native sample of *P. leniusculus* (117 native records) could by itself explain the observed patterns. *P. clarkii* was used as a benchmark because it had a large native sample and a strong climate-dominated pattern. Starting from the full *P. clarkii* dataset, native records were subsampled without replacement from 2,208 to 117 across 100 repetitions. In each repetition, a random forest was fitted and the thematic importance profile and top-10 variable overlap with the full-data model were recorded (Table S4; Figure S8).

Pseudo-absence sensitivity analysis for the separate niche models compared the original strategy (pseudo-absences from other species with the same status) with an alternative strategy using all non-focal crayfish records regardless of status. This tested whether the niche reorganisation conclusions were dependent on background definition (Table S3; Figure S7).

## 3 Results

### 3.1 Classification performance

Decision tree classifiers achieved high cross-validated performance in distinguishing native from invasive occurrences for all five species (Table 2). Classification accuracy ranged from 0.965 (*F. rusticus*) to 0.999 (*F. limosus*), and ROC AUC ranged from 0.955 to 0.998.

**Table 2.**
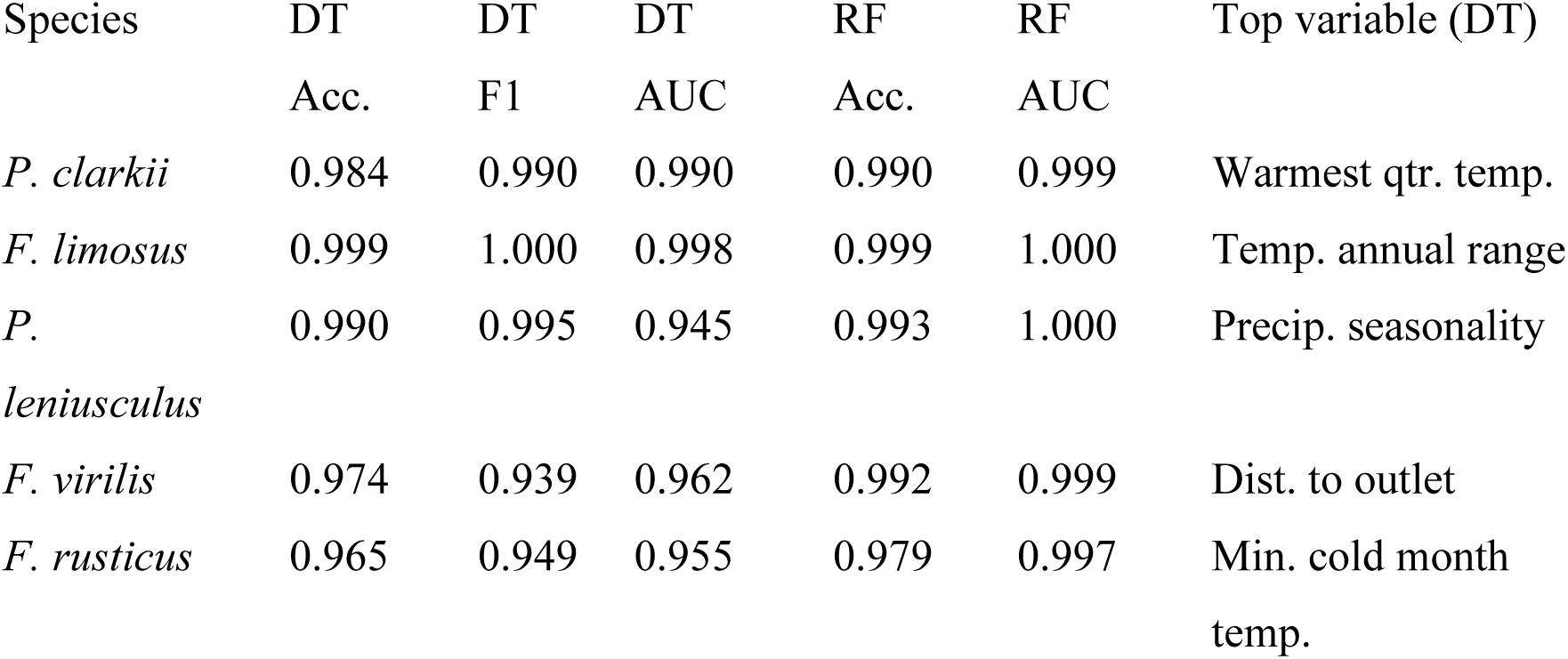
Decision tree and random forest classification performance for distinguishing native from invasive occurrences.

| Species | DT<br>Acc. | DT<br>F1 | DT<br>AUC | RF<br>Acc. | RF<br>AUC | Top variable (DT) |
| --- | --- | --- | --- | --- | --- | --- |
| *P. clarkii* | 0.984 | 0.990 | 0.990 | 0.990 | 0.999 | Warmest qtr. temp. |
| *F. limosus* | 0.999 | 1.000 | 0.998 | 0.999 | 1.000 | Temp. annual range |
| *P. leniusculus* | 0.990 | 0.995 | 0.945 | 0.993 | 1.000 | Precip. seasonality |
| *F. virilis* | 0.974 | 0.939 | 0.962 | 0.992 | 0.999 | Dist. to outlet |
| *F. rusticus* | 0.965 | 0.949 | 0.955 | 0.979 | 0.997 | Min. cold month temp. |

*Faxonius limosus* showed the highest separability, with a tree of only depth 4 and 8 leaves achieving near-perfect discrimination, indicating that its native and invasive ranges occupy almost entirely non-overlapping environmental spaces. The within-continent invaders (*F. virilis* and *F. rusticus*) showed slightly lower but still very high classification performance, consistent with greater environmental overlap between native and invaded ranges within the same continent.

Random forest models confirmed these results, with cross-validated accuracies of 0.979–0.999 and ROC AUC of 0.997–1.000 across all species. Specifically, RF accuracy/AUC were 0.990/0.999 for *P. clarkii*, 0.999/1.000 for *F. limosus*, 0.993/1.000 for *P. leniusculus*, 0.992/0.999 for *F. virilis*, and 0.979/0.997 for *F. rusticus*. Depth sensitivity analysis (depths 3–8) confirmed that decision tree performance was robust to tree complexity: accuracy gains from depth 3 to 5 were modest (1–2 percentage points), while increasing depth from 5 to 8 yielded negligible improvement (<0.5 percentage points). Notably, the high classification performance extended to within-continent species whose native and invasive ranges are not separated by ocean barriers, indicating that the environmental signal captured by the models is not merely a proxy for continental-scale geographic separation. The top-ranking features were stable across all depths tested (Figure S9).

### 3.2 The invasion pathway dichotomy

Aggregation of feature importances by environmental variable type revealed a striking dichotomy between invasion pathway types (Figure 1). For all three intercontinental invaders, climatic variables accounted for 91–93% of total decision tree feature importance: 93% for *P. clarkii* (dominated by mean temperature of the warmest quarter, l_CLI38, with 65.7% importance), 91% for *F. limosus* (temperature annual range, l_CLI25, 89.7%), and 92% for *P. leniusculus* (precipitation seasonality, l_CLI58, 75.4%). Topographic variables contributed only 5–9% for these species, and soil and land cover were negligible.

**Figure 1.**
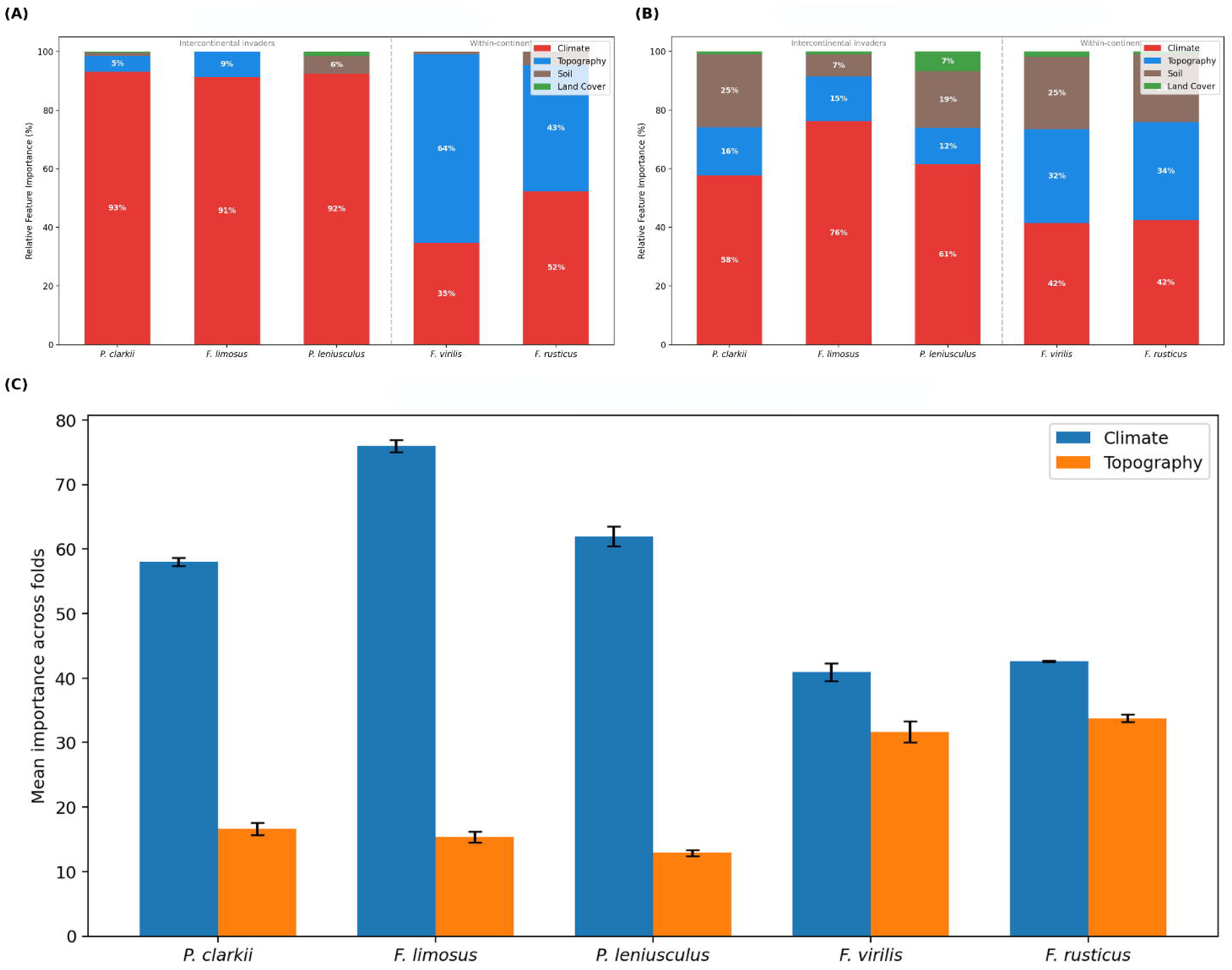
The invasion pathway dichotomy in environmental drivers of niche shift. (A) Relative feature importance aggregated by environmental variable type (Climate, Topography, Soil, Land Cover) from decision tree classifiers (depth = 5) for each species. The dashed line separates intercontinental invaders (left) from within-continent invaders (right). (B) Corresponding importance profile from random forest models (500 trees, Gini importance). (C) Cross-validation stability of climate and topography importance across 5-fold random forest cross-validation, showing mean ± SD. Error bars confirm that the dichotomy is stable across data partitions.

In contrast, the two within-continent invaders showed a fundamentally different importance profile. For *F. virilis*, topographic variables dominated at 64% of total importance, with the single most important variable being distance to basin outlet (l_TOP30, 62.7%), a network-position metric that reflects longitudinal position within the river network and is strongly associated with connectivity, dispersal pathways, stream size, and habitat heterogeneity, all of which are known to structure crayfish distributions (Reynolds et al., 2013). These variable captures where within the river network the species occurs, rather than what climate it experiences. Climate contributed only 35%. For *F. rusticus*, the balance was more even: climate at 52% and topography at 43%, with the top two features being minimum temperature of the coldest month (l_CLI21, 42.9%) and topological stream dimension (l_TOP109, 42.1%), a metric reflecting the branching complexity of the upstream network and thus the degree of hydrological connectivity available to the population. Soil and land cover variables contributed minimally across all species (<5%).

Random forest analysis broadly preserved the same ecological grouping (Figure 1B). Under Gini importance, climate accounted for 57.7% of total importance in *P. clarkii*, 76.2% in *F. limosus*, and 61.5% in *P. leniusculus*, whereas topography contributed 16.4%, 15.3%, and 12.4%, respectively. SHAP-based summaries yielded the same qualitative pattern: climate 61.8%, 75.8%, and 56.0% for the three intercontinental species; topography 15.5%, 14.2%, and 13.6%. For the within-continent invaders, RF Gini importance gave 41.5% climate and 31.9% topography for *F. virilis*, and 42.4% climate and 33.5% topography for *F. rusticus*; SHAP values were closely similar (39.0/32.9% and 43.8/33.6%, respectively). The redistribution of importance into soil and topography under random forests (particularly for *P. clarkii*, where soil reached 24.7%) reflects the ensemble’s ability to capture secondary environmental structure that the sparse decision tree solutions omit, but the central climate-versus-topography dichotomy was fully preserved (Figure S1). Decision tree feature importance rankings for all species are shown in Figure S10. This redistribution likely reflects the ensemble’s capacity to capture secondary, correlated environmental gradients, such as soil properties that covary with climatic regimes, that are not selected by the greedy, single-path structure of individual decision trees. Crucially, the redistribution did not alter the dominant ranking of climate versus topography across invasion pathways, confirming that the ecological pattern is robust to model architecture.

At the individual-variable level, the top-ranked features were ecologically coherent across methods (Figure 2). In *P. clarkii*, the dominant RF variables were climatic: mean temperature of the warmest quarter (l_CLI38), annual mean air temperature (l_CLI2), and precipitation seasonality (l_CLI57). In *F. limosus*, the top variables were temperature annual range (l_CLI25), temperature seasonality (l_CLI13), and warm-quarter precipitation. In *P. leniusculus*, precipitation seasonality (l_CLI58), precipitation of the warmest quarter (l_CLI69), and other precipitation-temperature combinations dominated. By contrast, *F. virilis* and *F. rusticus* both showed major contributions from network/topographic variables such as distance to basin outlet (l_TOP30) and topological dimension of streams (l_TOP109), together with climatic variables. Local-scale (90 m segment) variables dominated over upstream (catchment-aggregated) variables for all species (89–100% of importance), though *F. rusticus* showed the highest upstream contribution (∼11%).

**Figure 2.**
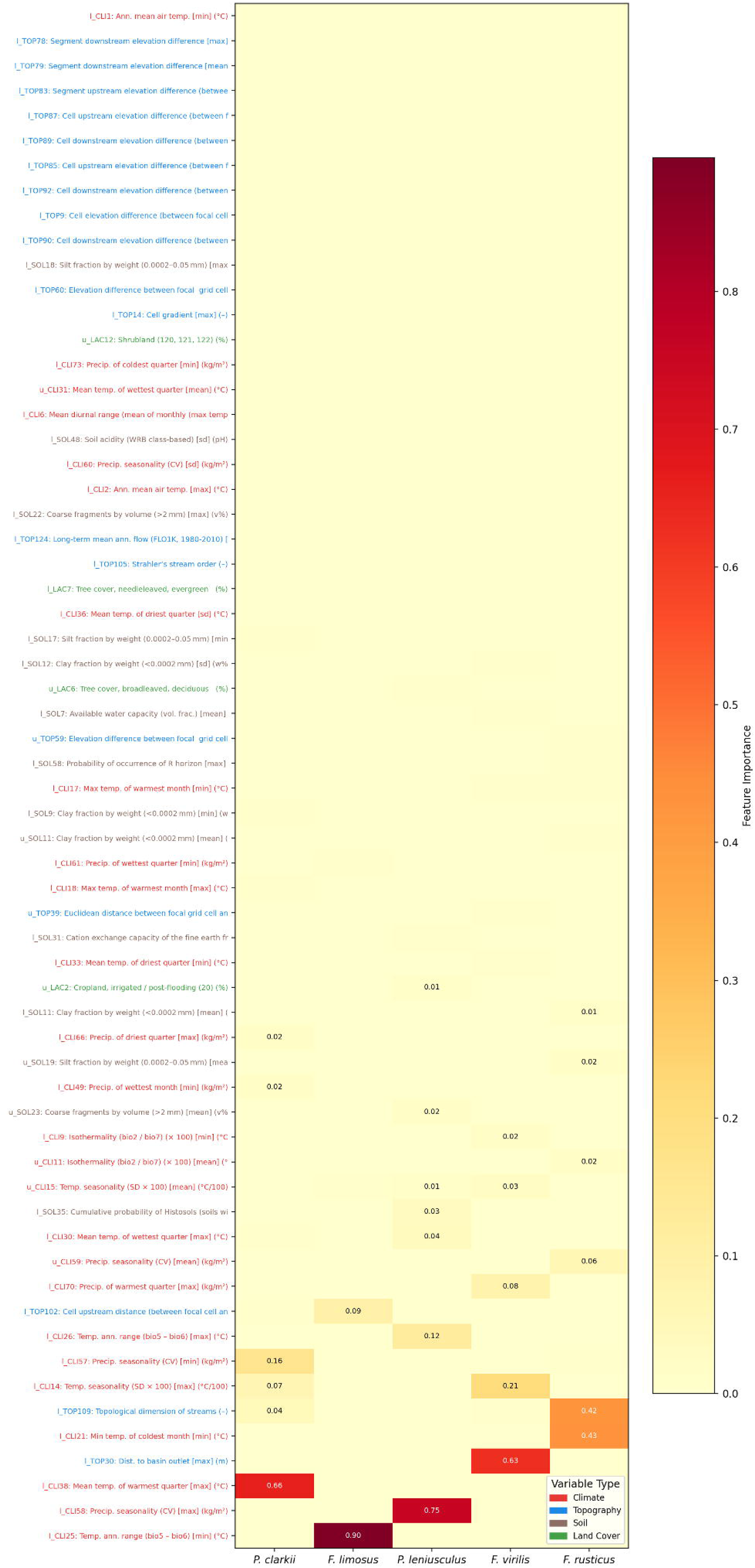
Feature importance heatmap across species. Rows represent individual environmental variables (top features from each species’ decision tree); columns represent species. Colour intensity indicates Gini importance. Variable labels include the variable code, full name, and thematic domain (colour-coded: red = Climate, blue = Topography, brown = Soil, green = Land Cover).

### 3.3 Robustness and statistical validation

The permutation null model supported the reality of the intercontinental–within-continent dichotomy when the contrast was expressed as the difference between climate and topography importance. Using random forest Gini importance, the observed between-group contrast was 41.20, whereas the permutation distribution was centred close to zero (mean = −0.75, SD = 21.80), yielding an empirical P = 0.001 for both one-sided and two-sided tests. The SHAP-based analysis produced an almost identical result: observed contrast 41.95, null mean = −0.63, SD = 22.28, P = 0.001. The ratio-based statistic was weaker (P = 0.100 for both Gini and SHAP), indicating that the signal is better captured by an additive contrast, probably because ratios become unstable when topographic importance is very small in strongly climate-dominated species (Table S1; Figure S6).

Cross-validation fold stability was high for all species. Standard deviations of theme-level importance across folds were ≤1.6%: *P. clarkii* showed 58.0 ± 0.7% climate and 16.6 ± 1.0% topography; *F. limosus* showed 75.9 ± 1.0% climate and 15.3 ± 0.8% topography; *P. leniusculus* showed 62.0 ± 1.5% climate and 12.9 ± 0.5% topography (Figure 1C). For the within-continent species, *F. virilis* had 40.9 ± 1.3% climate and 31.7 ± 1.6% topography; *F. rusticus* had 42.6 ± 0.1% climate and 33.8 ± 0.6% topography. The low standard deviations confirm that the dichotomy is not driven by a single data partition. At the individual-variable level, the top predictors recurred across folds for all species: in *P. clarkii*, l_CLI38, l_CLI2, l_TOP102, l_CLI18, and l_CLI57 appeared consistently among the highest-ranked variables; in *F. limosus*, l_CLI25, l_CLI13, l_CLI69, and l_TOP102 were stable across folds (Table S2; Figures S2–S3).

The sample-size sensitivity analysis confirmed that the climatic signal was not an artefact of small native sample size. When *P. clarkii* native records were downsampled from 2,208 to 117 (matching *P. leniusculus*), the resulting models still performed extremely well (mean accuracy 0.995 ± 0.001, ROC AUC 0.996 ± 0.003), and the thematic importance profile remained very similar to the full-data benchmark: 54.9 ± 2.2% climate, 17.5 ± 1.7% topography, 26.7 ± 2.4% soil, 0.9 ± 0.2% land cover (versus 58.5%, 15.9%, 24.7%, 0.9% in the full model). At the variable level, l_CLI18, l_CLI2, l_CLI38, and l_CLI57 appeared in the top 10 in all 100 repetitions, and the mean overlap between subsampled and full-data top-10 variables was 8.75 out of 10. This result indicates that the climate-dominated pattern observed in *P. leniusculus* is unlikely to be a trivial artefact of limited native sampling.

### 3.4 Classical overlap metrics versus machine learning

Classical niche overlap metrics showed that native and invasive environmental spaces overlapped to varying degrees across species (Figure 3A). Schoener’s D ranged from 0.299 (*F. rusticus*) to 0.619 (*P. clarkii*), and Warren’s I from 0.565 to 0.859. The highest overlap was observed in *P. clarkii* (D = 0.619, I = 0.859), followed by *F. limosus* (D = 0.480, I = 0.757), *P. leniusculus* (D = 0.369, I = 0.603), *F. virilis* (D = 0.326, I = 0.608), and *F. rusticus* (D = 0.299, I = 0.565).

**Figure 3.**
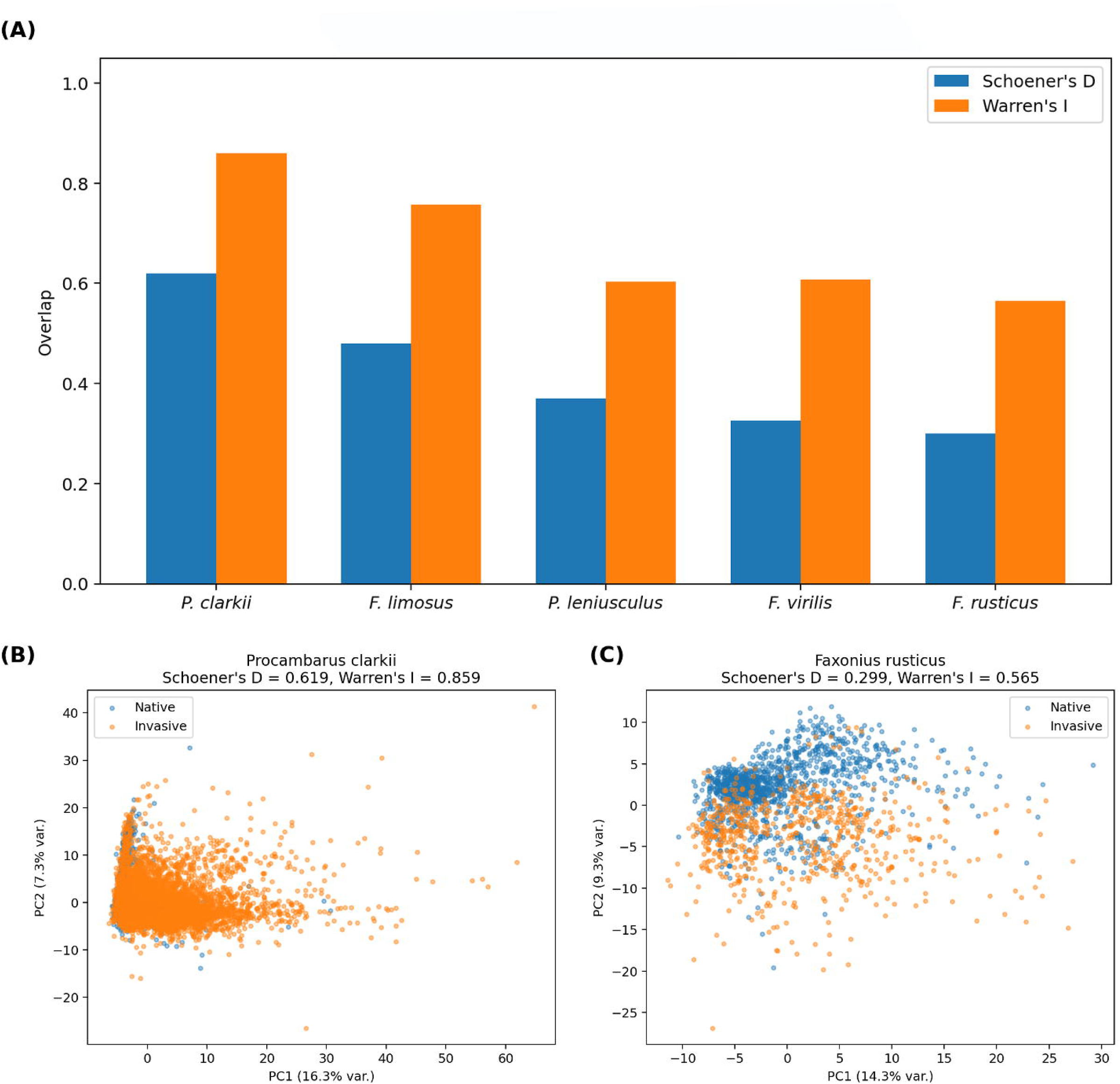
Classical niche overlap metrics versus machine learning. (A) Schoener’s D and Warren’s I between native and invasive environmental spaces for each species. (B) PCA scatter plot for *P. clarkii* (intercontinental; highest overlap, D = 0.619), showing that high scalar overlap coexists with strongly climate-dominated niche reorganisation. (C) PCA scatter plot for *F. rusticus* (within-continent; lowest overlap, D = 0.299), showing that low overlap reflects topographic and network-position shifts rather than climatic shifts.

Critically, these scalar metrics did not recover the invasion pathway dichotomy identified by the machine learning analyses. The overlap values yielded a gradient from relatively high overlap (*P. clarkii*) to lower overlap (*F. rusticus*), but they did not identify the qualitative difference between intercontinental invaders, whose native-versus-invasive separation is mainly climatic, and within-continent invaders, whose separation includes a much stronger topographic component. *Procambarus clarkii* had the highest classical overlap, yet still showed a strongly climate-dominated native-versus-invasive contrast under both decision tree and random forest analysis (Figure 3B). Conversely, the lower-overlap within-continent species did not simply differ by ‘more shift’ in a scalar sense; they differed in the *type* of environmental structure underlying that shift (Figure 3C). We note that PCA-based overlap metrics are sensitive to variable scaling and the choice of included variables, which may affect the absolute values of D and I; however, the relative ranking across species and the disconnect with axis-specific patterns are robust to these choices. D and I collapse niche difference into a single number, whereas the interpretable machine learning approach identifies *which* environmental axes are being reorganised (PCA scatter plots and kernel density maps for all five species are provided in Figures S4 and S5).

### 3.5 Niche reorganisation versus threshold shift

Separate niche models trained independently on native-range and invasive-range data revealed that most species are governed by entirely different environmental axes in their two ranges. *Procambarus clarkii* and *F. limosus* did not share any top splitting variables between their native and invasive niche models under the chosen model structure, suggesting substantial niche reorganisation (Table S6). This absence should be interpreted cautiously, as alternative but correlated variables may represent similar environmental gradients; nevertheless, the lack of overlap in the selected predictors indicates that the dominant axes of ecological differentiation change between ranges.

For *P. clarkii*, the native niche model identified warmest-quarter temperature (>26.7°C), diurnal range, and soil pH as the primary determinants, while the invasive niche model relied on annual mean temperature (>12.4°C), precipitation seasonality, and maximum warmest-month temperature. In its native range, *P. clarkii* is distinguished from co-occurring species as a warm-specialist among warm-adapted species; in its invasive range, it is simply the warm-tolerant species in a cooler community, a qualitative shift in niche identity.

*Pacifastacus leniusculus* showed one shared variable between range models: temperature seasonality, with a threshold shift of +46.8% (from 533 to 783 °C/100; Figure 5). This indicates that signal crayfish in Europe tolerate substantially more continental climates than in their native Pacific Northwest range.

The within-continent invaders showed more niche continuity. *Faxonius rusticus* had three shared variables: temperature annual range (shift: +4.7%), maximum warmest-month temperature (shift: −10.0%, from 30.6 to 27.6°C), and temperature seasonality (shift: −2.7%). This species is expanding into areas with slightly cooler summers but similar overall thermal regimes. *Faxonius virilis* had one shared variable (temperature annual range), with a −10.0% shift indicating invasion of areas with less extreme annual temperature variation.

The pseudo-absence sensitivity analysis confirmed that native-range niche models were generally robust to background definition, with climate remaining dominant under both strategies across species. Invasive-range models were more sensitive: for example, *F. limosus* invasive shifted from 67.6% topography under the original strategy to 83.6% under the alternative, and *P. leniusculus* invasive showed a substantial shift in the relative contribution of soil versus climate. These results indicate that separate niche models provide valuable but context-dependent insight into niche reorganisation, and should be interpreted more cautiously than the primary native-versus-invasive classification (Table S3; Figure S7). *[Figure 4 here]*

**Figure 4.**
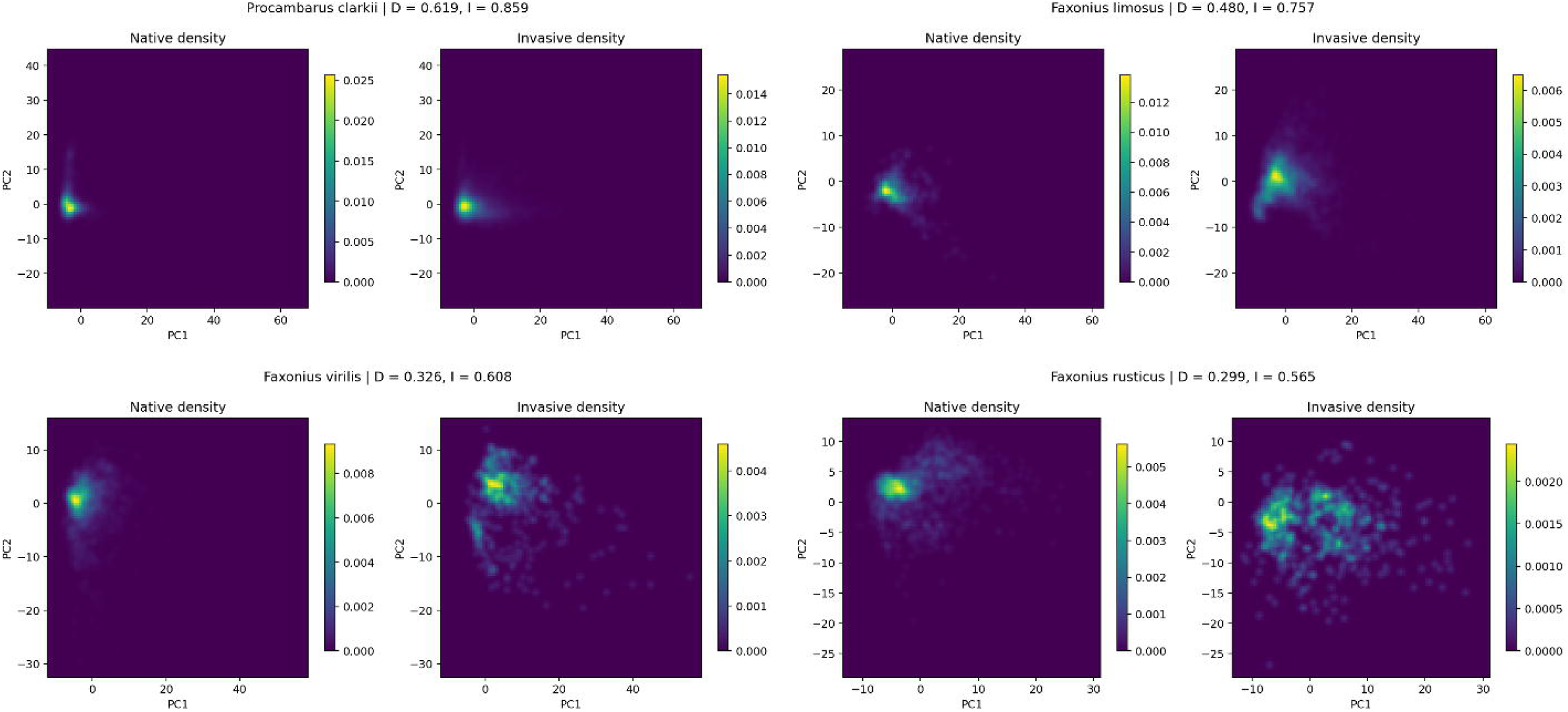
Niche reorganisation in environmental space. Kernel density maps of native (left) and invasive (right) occurrences in PCA space for (A) *P. clarkii*, (B) *F. limosus*, (C) *F. virilis*, (D) *F. rusticus*. Values of Schoener’s D and Warren’s I are shown above each panel. Intercontinental invaders (A, B) show displacement of the density centre along PC1 (climate-loaded); within-continent invaders (C, D) show partial overlap with redistribution of density across both axes (all five species shown in Figure S5).

**Figure 5.**
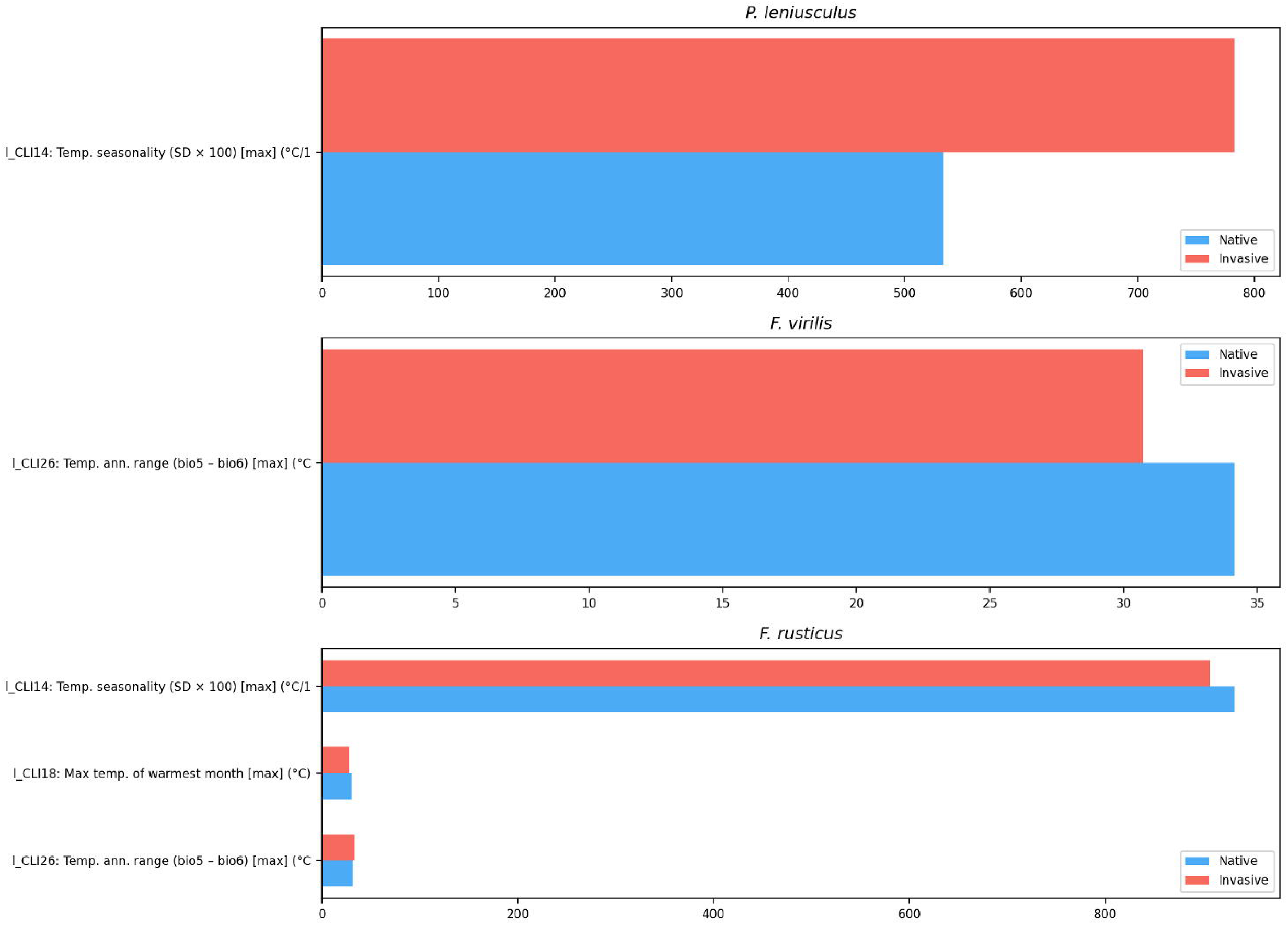
Niche threshold comparison for species with shared variables between native and invasive niche models. Horizontal bars show the threshold value of the shared variable in the native model (blue) and invasive model (red). *P. leniusculus* shows the largest shift: +46.8% in temperature seasonality between native (Pacific Northwest) and invasive (European) ranges.

## 4 Discussion

### 4.1 Network-aware variables enable high-resolution niche discrimination

The high classification accuracies achieved in this study (96.5–99.9% for decision trees, 97.9–99.9% for random forests) demonstrate that the native and invasive ranges of all five study species occupy detectably different environmental spaces when characterised using network-aware freshwater variables. This result validates the analytical utility of the GeoTraits database and supports the broader argument that hydrologically resolved environmental descriptors capture ecological variation that terrestrial climate grids systematically miss (Domisch et al., 2015; Cid et al., 2022). The near-perfect separability of *F. limosus* (99.9% accuracy with a tree of only depth 4) is particularly notable, indicating that eastern North American and European freshwater environments are so environmentally distinct, at least along the axes captured by the Hydrography90m and CHELSA datasets, that a handful of threshold rules suffice to discriminate the two ranges. For the within-continent invaders, where climatic differences between native and invaded ranges are subtle, the inclusion of topographic and network-position variables proved critical: *F. virilis* and *F. rusticus* could not have been separated at these accuracies using climate variables alone.

### 4.2 The invasion pathway dichotomy: climate versus landscape structure

The most striking finding of this study is the qualitative difference in the environmental dimensions driving niche reorganisation between intercontinental and within-continent invaders. For species that crossed oceans (*P. clarkii*, *F. limosus*, *P. leniusculus*), the invasive range is distinguished from the native range almost entirely by climatic variables, with 91–93% of decision tree importance and 58–76% of random forest importance falling in the Climate domain. This pattern is consistent with macroecological expectation: intercontinental dispersal, mediated by human translocation, places organisms in fundamentally different thermal and precipitation regimes, and the primary niche axis along which adjustment occurs is the one that varies most between continents. The specific climatic discriminators differ by species in ecologically interpretable ways: *P. clarkii* invasions are governed by warmest-quarter temperature (reflecting the species’ subtropical origin and its establishment in cooler European climates), *F. limosus* by temperature annual range (reflecting the shift from maritime-influenced eastern North America to more continental European climates), and *P. leniusculus* by precipitation seasonality (reflecting the shift from the wet Pacific Northwest to drier continental European climates).

For the within-continent invaders (*F. virilis*, *F. rusticus*), the climate is broadly similar between native and invaded ranges, both lie within temperate North America, and the distinguishing axes shift to topography and network position. The dominance of distance to basin outlet for *F. virilis* (62.7% of total importance) is particularly revealing: this metric reflects where within the river network a species occurs, not what climate it experiences.

*Faxonius virilis* invasive populations tend to establish in different positions along the river continuum compared to native populations, suggesting colonisation of structurally different habitats (e.g., different stream orders, different connectivity profiles) rather than climatically different regions. For *F. rusticus*, the co-dominance of minimum coldest-month temperature and topological stream dimension indicates a combined thermal and structural niche shift consistent with the species’ known expansion from the Ohio River basin into larger, more connected waterways of the Great Lakes region.

The robustness of this dichotomy across analytical approaches, confirmed by both decision trees and random forests, both Gini and SHAP importance measures, stable across cross-validation folds with standard deviations ≤1.6%, and significant under permutation testing (P = 0.001 for the additive contrast), indicates that this is a genuine ecological pattern rather than a methodological artefact. The limited power of the ratio-based permutation test (P = 0.100) does not undermine this conclusion; it simply reflects the instability of ratios when the denominator (topographic importance) is very small in climate-dominated species.

### 4.3 Niche reorganisation, not niche shift

The separate niche model analysis revealed a more fundamental pattern than simple threshold shifts along conserved axes. For *P. clarkii* and *F. limosus*, the environmental rules governing occurrence changed entirely between ranges, zero shared variables between native and invasive niche models. This goes beyond the niche shift/conservatism framework as traditionally conceived (Broennimann et al., 2007; Petitpierre et al., 2012; Guisan et al., 2014) by demonstrating that invasion involves not merely a displacement along existing niche axes but a reorganisation of which axes matter.

The case of *P. clarkii* illustrates this most vividly. In its native range, the species is distinguished from co-occurring crayfish by warmest-quarter temperature, diurnal range, and soil pH, variables that differentiate a warm-specialist within a warm-adapted community. In its invasive range, it is distinguished by annual mean temperature, precipitation seasonality, and maximum warmest-month temperature, variables that identify a warm-tolerant generalist within a cool-adapted community. The species’ fundamental physiology has presumably not changed, but the environmental axes that matter for predicting its occurrence have shifted entirely because the competitive and climatic context is different. This represents a qualitative change in niche identity, not merely a quantitative displacement.

The +46.8% threshold shift in temperature seasonality for *P. leniusculus* is also noteworthy: signal crayfish in Europe tolerate climates with substantially greater temperature variation than in their native Pacific Northwest range, consistent with their successful establishment across continental Europe from Scandinavia to the Mediterranean. The within-continent species, by contrast, showed more niche continuity, consistent with the expectation that intra-continental range expansion involves less dramatic environmental novelty.

This finding has direct implications for invasion risk modelling. Species distribution models trained on native-range environmental associations may fail to predict invasive ranges not because the species’ fundamental niche has changed, but because the relevant niche axes shift when the competitive and environmental context changes. Transferability of SDMs across ranges (Petitpierre et al., 2012) may be limited not by niche evolution but by context-dependent niche expression, a distinction with profound consequences for forecasting future invasion risk.

### 4.4 The limits of scalar overlap metrics

Our comparison of classical niche overlap metrics with interpretable machine learning reveals a critical blind spot in scalar overlap summaries. Schoener’s D and Warren’s I provided a gradient from relatively high overlap (*P. clarkii*, D = 0.619) to lower overlap (*F. rusticus*, D = 0.299), but this gradient bore no systematic relationship to the qualitative nature of niche reorganisation. *Procambarus clarkii* had the highest classical overlap yet showed the most purely climate-dominated niche reorganisation under both decision tree and random forest analysis. Conversely, the lower-overlap within-continent species did not simply show ‘more shift’ in a scalar sense; they differed fundamentally in the *type* of environmental structure underlying that shift.

This disconnect arises because D and I are computed in reduced environmental space (typically PCA) and summarise total divergence without attributing it to specific environmental axes. The machine learning approach, by contrast, identifies exactly which variables drive the separation and how much each contributes. For invasion ecology, this distinction is not merely methodological but substantive: a manager who knows that a species’ invasive range is distinguished by climate can direct monitoring towards regions projected to enter the species’ thermal envelope, whereas one who knows that the distinction is topographic can focus on connectivity and network position. Scalar overlap metrics cannot inform such decisions.

This does not render classical overlap metrics obsolete. They remain valid summaries of total niche divergence and are useful for broad comparative analyses across many species (Petitpierre et al., 2012). But for mechanistic understanding of invasion, for asking not just *how different* a species’ niche is in its invasive range, but *how it is different*, interpretable machine learning applied to richly resolved environmental data offers a complementary and, for mechanistic questions, qualitatively richer answer.

### 4.5 Methodological considerations and future directions

Several limitations should be acknowledged. First, the pseudo-absence sensitivity analysis revealed that separate niche models (Analysis 2) were not uniformly robust to background definition, particularly for invasive-range models. Native-range models were generally stable, with climate remaining dominant across strategies, but invasive-range models showed greater sensitivity, for example, *F. limosus* invasive shifted from 67.6% to 83.6% topography depending on background definition. This is scientifically informative in itself: it shows that the invasive-range niche signal is more background-dependent than the primary native-versus-invasive signal, presumably because invasive ranges are characterised relative to a less well-defined ecological context. We recommend that separate niche models be interpreted as valuable secondary evidence rather than definitive assessments (Table S3; Figure S7).

Second, the unit of inference for the pathway dichotomy is the species (n = 5), and the permutation null model has inherently limited power at this sample size. The significant result for the additive contrast (P = 0.001) is encouraging, Accordingly, the dichotomy identified here should be interpreted as a pattern emerging across a small set of species-level replicates, and thus as a hypothesis requiring broader taxonomic validation. The GeoTraits database is growing, and several additional species (e.g., *Cherax destructor*, *Procambarus virginalis*) are approaching the sample-size thresholds used here.

Third, spatial autocorrelation inherent in species occurrence data may inflate classification performance, because geographically proximate records share environmental similarity independently of species’ ecological requirements. We did not implement spatial blocking or spatial cross-validation, which could provide a more conservative estimate of true ecological separability. Several features of our design provide partial, though not definitive, reassurance: deduplication to one record per stream segment reduces fine-scale spatial clustering; the within-continent species, whose native and invasive ranges overlap geographically within temperate North America, were still classified at >96% accuracy, indicating that the signal is not purely a continental proxy; and the consistency of results across decision trees and random forests suggests that the patterns reflect broad environmental structure rather than local spatial artefacts. These results should therefore be interpreted conservatively until confirmed with spatially explicit validation. Future analyses should incorporate spatially stratified cross-validation to quantify the contribution of spatial structure to classification performance.

More broadly, the classification models quantify environmental differentiation conditional on the observed geographic separation of native and invasive ranges, rather than providing a fully geography-independent estimate of niche divergence. Fourth, the analysis is based on static, long-term environmental averages and does not capture the temporal dynamics of invasion. Future work should integrate temporal environmental trajectories to assess whether niche reorganisation is ongoing, has stabilised, or is reversing as invasive populations adapt.

Fifth, sample sizes vary substantially across species and ranges, from 117 native records for *P. leniusculus* to 8,527 invasive records for *P. clarkii*. The sample-size sensitivity analysis demonstrated that the climatic signal is preserved even under drastic subsampling (mean top-10 overlap of 8.75/10 at n = 117), but data-poor species will inevitably have less precisely characterised niches (Table S4).

Finally, while decision trees offer unmatched interpretability, they capture greedy, axis-aligned partitions of predictor space. The consistent confirmation of all major patterns by 500-tree random forests with both Gini and SHAP importance substantially mitigates this concern. Future analyses could extend this framework with gradient-boosted trees and partial dependence plots to capture more nuanced predictor interactions.

## 5 Conclusions

This study indicates that invasive freshwater crayfish undergo niche reorganisation, not merely niche shift, upon invasion, and that the axis of this reorganisation is consistently associated with invasion pathway type. Intercontinental invaders, translocated across oceans by human agency, reorganise their ecological niches along climatic axes, reflecting establishment in fundamentally different thermal and precipitation regimes. Within-continent invaders, expanding through connected or adjacent waterways, reorganise along topographic and network-position axes, colonising structurally different positions within similar-climate river networks. This dichotomy, robust across classification methods, importance metrics, and cross-validation partitions, reveals that the traditional niche shift/conservatism framework, which asks how much niches change, misses a fundamental ecological question: what changes. Classical niche overlap metrics, while useful summaries of total divergence, are blind to this qualitative distinction. By combining the first globally comprehensive, network-aware freshwater environmental database (GeoTraits) with interpretable machine learning, we show that the nature of niche reorganisation during invasion is structured and consistently associated with invasion pathway type. These findings provide an analytical foundation for mechanistically informed invasion risk assessment: knowing not only that a species’ niche will change upon invasion, but along which environmental axes it will change, is essential for directing monitoring, forecasting spread, and prioritising management in freshwater ecosystems.

## Data Availability Statement

The Global Crayfish Database of Geospatial Traits is freely available through the World of Crayfish® platform (https://world.crayfish.ro/). Analytical code and processed datasets supporting this study are available at https://anonymous.4open.science/r/niche-shift-4285 (anonymised for blind review). The underlying dataset is archived at Mendeley Data [DOI removed for review].

## Supplementary Figures

Figure S1. Comparison of relative feature importance aggregated by environmental variable type between decision tree classifiers (depth = 5; light bars) and random forest models (500 trees; dark bars) for each of the five study species. Four panels show Climate, Topography, Soil, and Land Cover separately.

Figure S2. Variable-type importance stability (5-fold CV) for each species. Bar height represents mean importance across folds; error bars represent ± 1 SD. (A) P. clarkii, (B) F. limosus, (C) P. leniusculus, (D) F. virilis, (E) F. rusticus.

Figure S3. Top variable stability (5-fold CV) for each species. Horizontal bars show mean feature importance across folds for the 12 highest-ranked variables; error bars show ± 1 SD. (A) P. clarkii, (B) F. limosus, (C) P. leniusculus, (D) F. virilis, (E) F. rusticus.

Figure S4. PCA scatter plots of native (blue) and invasive (orange) occurrences in two-dimensional environmental space for all five species: (A) P. clarkii (D = 0.619, I = 0.859), (B) F. limosus (D = 0.480, I = 0.757), (C) P. leniusculus (D = 0.369, I = 0.603), (D) F. virilis (D = 0.326, I = 0.608), (E) F. rusticus (D = 0.299, I = 0.565).

Figure S5. Kernel density maps of native (left) and invasive (right) occurrences in PCA space for all five species: (A) P. clarkii, (B) F. limosus, (C) P. leniusculus, (D) F. virilis, (E) F. rusticus. Colour scale represents normalised occupancy density.

Figure S6. Null model permutation distributions. (A) RF Gini climate−topography contrast, (B) RF SHAP climate−topography contrast, (C) RF Gini climate/topography ratio, (D) RF SHAP climate/topography ratio. Dashed line indicates observed statistic.

Figure S7. Pseudo-absence sensitivity: strategy comparison bar plots showing thematic importance under Strategy A (other species with same status) and Strategy B (all non-focal species) for each species and range.

Figure S8. Sample-size sensitivity analysis. (A) Thematic importance stability under subsampling of P. clarkii native records to n = 117 (100 iterations). (B) Top-variable frequency across iterations.

Figure S9. Decision tree classifiers (depth = 5) for each species: (A) P. clarkii, (B) F. limosus, (C) P. leniusculus, (D) F. virilis, (E) F. rusticus. Node colours indicate predicted class (blue = native, orange = invasive).

Figure S10. Top 20 environmental variables (decision tree Gini importance) distinguishing native from invasive occurrences for each species: (A) P. clarkii, (B) F. limosus, (C) P. leniusculus, (D) F. virilis, (E) F. rusticus.

Figure S11. SHAP summary plots from random forest models for each species: (A) P. clarkii, (B) F. limosus, (C) P. leniusculus, (D) F. virilis, (E) F. rusticus. Each dot represents a record; position on the x-axis shows the SHAP value; colour indicates feature value (red = high, blue = low).

## Supplementary Tables

Table S1. Null model permutation test results for the intercontinental versus within-continent dichotomy. The observed statistic is the between-group contrast (intercontinental mean minus within-continent mean). P-values are empirical with +1 correction (999 permutations).

Table S2. Theme-level variable importance stability across 5-fold cross-validation (random forest, 500 trees). Values are mean ± SD of total importance (%) across folds.

Table S3. Pseudo-absence sensitivity analysis for separate niche models. (A) Native-range models. (B) Invasive-range models. Strategy A: pseudo-absences from other species with same status. Strategy B: all non-focal species regardless of status.

Table S4. Sample-size sensitivity analysis. P. clarkii native records subsampled from 2,208 to 117 across 100 iterations (random forest, 500 trees).

Table S5. Classical niche overlap metrics between native and invasive environmental spaces, including PCA variance explained by PC1 and PC2.

Table S6. Separate niche model summary. (A) Top splitting variables, thresholds, and importance for native and invasive models per species. (B) Threshold shifts for shared variables between native and invasive niche models.

Table S7. Decision tree feature importance (Gini importance) for all variables per species (Excel file with five sheets).

Table S8. Random forest Gini importance for all variables per species (Excel file with five sheets, 500 trees).

Table S9. Random forest SHAP importance (mean |SHAP|) for all variables per species (Excel file with five sheets, 500 trees).

## Notes

https://doi.org/10.17632/8j6mgp32fx.1

https://github.com/KristianMiok/niche-shift

### Competing Interest Statement

The authors have declared no competing interest.

